# Kin discrimination modifies fitness, spatial segregation and matrix sharing between strains with low relatedness in *Bacillus subtilis* biofilms

**DOI:** 10.1101/2021.09.24.461764

**Authors:** Maja Bolješić, Barbara Kraigher, Iztok Dogša, Barbara Jerič Kokelj, Ines Mandić Mulec

## Abstract

Microorganisms in nature form multicellular groups called biofilms. In biofilms bacteria embedded in a matrix of extracellular polymeric substances (EPS) interact intensely, due to their proximity to each other. Most studies have investigated genetically homogeneous biofilms, leaving a gap in knowledge on genetically heterogeneous biofilms. Recent insights show that a Gram-positive model bacterium, *Bacillus subtilis*, discriminates between strains of high (kin) and low (non-kin) phylogenetic relatedness, reflected in merging (kin) and boundaries (non-kin) between swarms. However, it is not clear how kinship between interacting strains affects their fitness, the genotype distribution, and the EPS sharing in floating biofilms (pellicles). To address this gap in knowledge we cultivate *B. subtilis* strains as mixtures of kin and non-kin strains in static cultures, allowing them to form pellicles. We show here that in non-kin pellicles only one strain’s fitness was reduced; at the same time, strains segregated into larger patches and exhibited decreased matrix sharing, as compared to kin and isogenic pellicles, in which both strains had comparable colony forming units (CFU) counts and more homogenous cell mixing. Overall, our results emphasize kin discrimination (KD) as a social behavior that shapes fitness, spatial segregation and sharing of the extracellular matrix in genetically heterogenous biofilms of *B. subtilis*.

**IMPORTANCE:** Biofilm communities have both beneficial and harmful effects on human societies in natural, medical and industrial environments. *Bacillus subtilis*, a Gram-positive and biotechnologically important bacterium, serves as a model for studying biofilms. Recent studies have shown that this species engages in kin discriminatory behavior during swarming, which may have implications for community assembly, thus being of fundamental importance. Effects of KD on fitness, genotype segregation and matrix sharing in biofilms is not well understood. By using environmental strains with determined kin types and integrated fluorescent reporters we provide evidence that KD involves antagonism of the dominant strain against non-kin, which has important implications for genotype segregation and sharing of matrix polysaccharides between producers and non-producers. Our results reveal novel consequences of KD and are important for advancing our fundamental understanding of microbial sociality, and its role in the assembly of multicellular groups and in the shaping of microbial diversity.

## Introduction

Biofilms are multicellular assemblages of bacteria that engage in intense social interactions that bring about positive (cooperative) or negative (antagonistic) fitness effects (1). Biofilms in natural habitats are the most dominant forms of microbial life and are predominantly composed of genetically diverse microorganisms (2–4). Nevertheless, most studies have focused on genetically homogenous biofilms.

The hallmark of biofilms is a matrix of self-generated extracellular polymeric substances (EPS) that glues cells together, mediates surface attachment, provides stability and improves survival in harsh environments (5). Matrix polymers serve as “public goods” because they are released and shared by cells joined in biofilm collectives (6). However, we have limited understanding of matrix sharing in genetically heterogeneous biofilms. According to Hamilton (7, 8), organisms apply kin discrimination (KD) to match their behaviors toward others according to their relatedness. Moreover, KD may limit the possibility of less related genotypes exploiting costly public goods during group living, due to cells’ sorting, avoidance or antagonism between non-kin (9, 10).

Studies on bacterial KD have mostly employed cooperative swarming as the test behavior to determine phenotypic responses of bacterial strains at the point of encounter, where a visible boundary between two strains was found to indicate non-kin interactions. In contrast, merging was found to be more common between swarms of very close kinship or between isogenic swarms (11–16). Recently, we showed that non-pathogenic, spore-forming, Gram-positive soil bacterium *Bacillus subtilis* engages in KD during swarming, where the frequency of the boundary appearance or merging at the meeting point of two swarms correlates with phylogenetic relatedness. Relatedness between interacting strains was defined by comparing the identity of four housekeeping genes (13) and by Average Nucleotide Identity (ANI) (17, 18) of orthologous gene pairs shared between two microbial genomes. Specifically, *B. subtilis* kin strains with 99.93 – 99.99% ANI and isogenic strains (“self” pairings, 100% ANI) exhibited merging behavior, whereas less related strains (98.73 – 98.84% ANI), which we also refer to as non-kin strains, formed clear boundaries (18). Moreover, Lyons *et al*. showed that less related *B. subtilis* strains also differ in a combination of KD loci, which comprise genes for antimicrobials, contact-dependent toxin–immunity pairs, and the enzymes involved in the synthesis of major matrix polysaccharide EpsA-O (16). However, we lack information on how KD affects fitness and cell sorting of different genotypes in biofilms (19).

*B. subtilis* is a model organism that is often used to investigate biofilm development on plant roots (13, 20, 21), on agar surfaces (22) or at the liquid–air interface, where *B. subtilis* constructs intricate pellicles (23–25). Pellicle formation is dependent on biofilm matrix polysaccharides EpsA-O and the TasA amyloid protein fibers, anchored through TapA to peptidoglycan (24, 26–28). Monocultures of *B. subtilis* matrix mutants form poor pellicles (26, 29, 30). In contrast, co-culturing the *tasA* and *EpsA-O* mutants that are isogenic in other loci compensates for the defect in pellicle formation (26, 28, 31). It is not known whether matrix sharing also occurs between phylogenetically different (non-kin) strains of *B. subtilis*.

We here hypothesize that low kinship will induce spatial segregation and decrease fitness of interacting *B. subtilis* strains in genetically mixed biofilm, where non-kin interactions between *B. subtilis* strains will limit matrix sharing between the producers and the Δ*epsA-O* mutant. To test these predictions we use selected *B. subtilis* strains isolated from one 1 cm^3^ soil sample (32) with previously determined genomic relatedness and KD phenotype (13, 16, 18).

Here, we provide evidence that two non-kin *B. subtilis* strains behave differently from kin or isogenic mixtures in pellicles. In non-kin pellicles the dominant strain decreases the fitness of the partner strain, whereas this dominance is not detected in kin pellicles. Also, in non-kin pellicles strain segregation is more prominent than in kin pellicles, where two strains remain more intermixed. Moreover, non-kin interactions also restrict sharing of EPS polysaccharides between the non-kin producer and the Δ*epsA-O* mutant in pellicles. The findings of this work make an important contribution to our understanding of how KD shapes bacterial biofilms, which is relevant for biofilm control and application.

## Results

### Kinship-dependent social interactions in *Bacillus subtilis* floating biofilm affect the fitness of less dominant strains

Bacteria in natural settings reside in genetically mixed collectives. To monitor the fitness of different strains in mixed biofilms (pellicles), we used strains, which originate from a microscale soil environment (32), and labeled them by constitutively expressed fluorescent reporters (yellow fluorescent protein-YFP or red fluorescent protein-mKate2) and associated antibiotic resistance (spectinomycin for strains marked with YFP, and chloramphenicol for strains marked with mKate2). These provided the means to determine the average fitness of each strain in pellicle competition assay by counting cells (CFU) on selective media. Two strains were inoculated together at a 1:1 ratio (10^5^ cells/ml each) in the biofilm-promoting minimal medium (MSgg) and then incubated under static conditions for 24 hours. The results showed that one strain gained a fitness advantage over the other in non-kin pellicles, whereas in kin pellicles both highly related strains reached similar final cell counts (**Fig. 1)**. Specifically, the average cell counts of the two kin strains in the pellicle (PS-216 mKate2 with PS-18 YFP or PS-68 YFP) did not change significantly. However, in non-kin pellicles PS-216 YFP had an average fitness advantage of 1.30-log CFU/ml (*P* < 0.001) and 1.58-log CFU/ml (*P* = 0.014) relative to PS-196 mKate2 or PS-218 mKate2, respectively. Similarly, PS-68 YFP had a significantly higher average fitness compared to non-kin PS-218 mKate2 (1.63-log CFU/ml; *P* = 0.024). In contrast, differentially labeled isogenic co-cultures (YFP and mKate2), which served as the control, showed no significant fitness modification (**Fig. S1**). Thus, these results are consistent with the prediction that KD-associated interactions lead to differential behavior of strains in non-kin biofilms, with one strain being dominant over the other.

**FIG 1.**
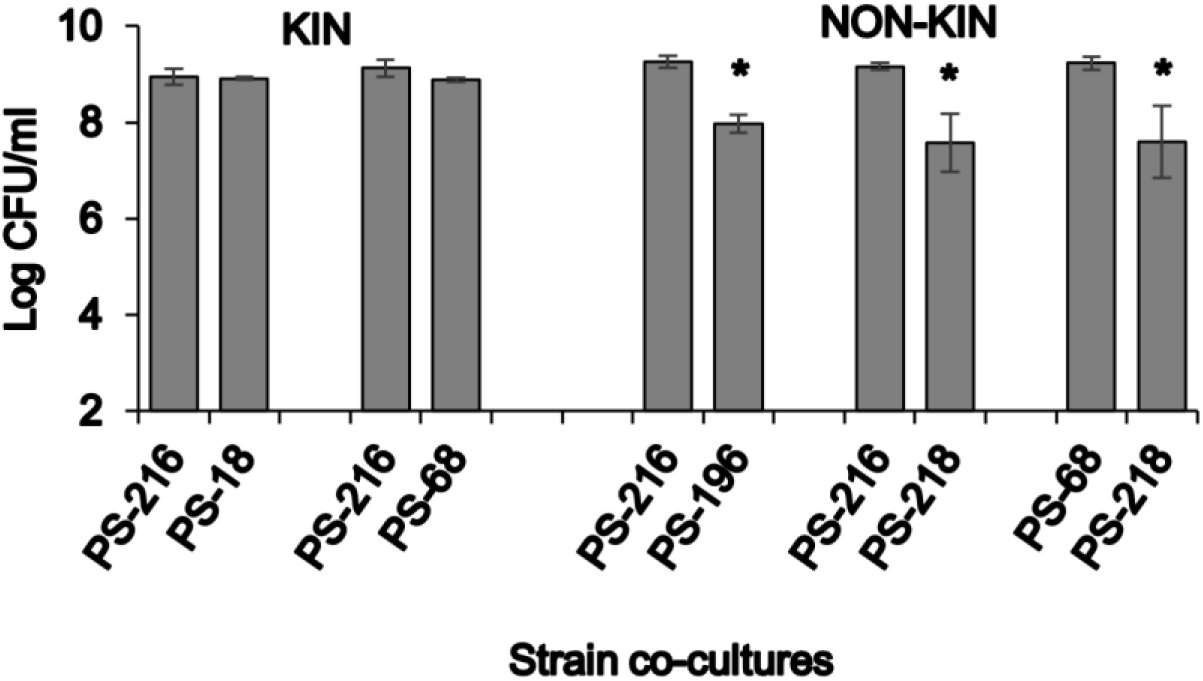
Cell frequency in pellicles of two kin and two non-kin strains. Two strains differentially labeled with different antibiotic resistance markers were grown in given combinations in 2 ml of MSgg for 24 hours at 37°C and the frequency of each strain was assessed in pellicles by CFU counts. Experiments were performed in at least three to four independent experimental replicates (n ≥ 3). *Indicates that the log CFU/ml is significantly different between the two strains in mixed pellicles (*t*-Student; two-tail * *P* < 0.05).

### KD affects spatial assortment of strains in biofilms

Given that during pellicle formation *B. subtilis* strains engage in interactions with differential fitness effects for kin and non-kin pairs, we predicted that these interactions may also impact their spatial distribution in pellicles. To test this prediction, we grew mixed cultures as described above using the same pairs of strains labeled with two different fluorescent markers (YFP and mKate2). We examined pellicles by confocal laser scanning microscopy (CLSM) after removing the liquid medium. By visually inspecting slices of confocal images of biofilm stacks, we concluded that isogenic and kin cells were more homogenously dispersed, whereas in non-kin pellicles one strain occupied a larger surface/area than the other and segregation seemed more pronounced (**Fig. 2** and **S2**). However, as the intensity of the fluorescent marker may have influenced our visual perception of the strain distribution in the images, we applied an additional unbiased estimator of cell segregation. Specifically, we developed multiscale spatial segregation analysis (MSSA) of the observed field of view (FOV) in the CLSM image biofilm’s stacks (detailed information available in Dogsa *et al*.; under consideration). Briefly, the FOV is a square with dimensions *d* x *d*, with *d* ranging from 12 μm to 1.2 mm. An example of such an analysis is given in **Fig. 3A**. When FOV is 1.2 mm, the segregation appears low. As we zoom in, and thereby decrease the size of FOV, the segregation becomes apparent and more intense in non-kin pellicles (**Fig. 3A and 3B**). For the experimental pellicle in the FOV of 12 μm x 12 μm the segregation level 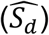 was 0.44 for the kin pellicle and 0.6 for the non-kin pellicle (**Fig. 3A**). If we assume that the overall ratio (in the whole microscopy image) of the two strains is 1:1, we find that at the lower size scale (FOV 12 μm x 12 μm) the ratio is always in favor of one strain. However, the effect is more pronounced in non-kin pairs (7:3 for kin vs. 12:3 for non-kin pair). This means that at the smallest FOV (12 μm x 12 μm) it is more likely that in the non-kin pellicle only one strain will occupy the whole 12 μm x 12 μm image, compared to the kin pellicle (**Fig. 3B**). To simplify the comparison of several segregation-level curves and several combinations of strains, we calculated the multiscale spatial segregation level (MSSL), which is 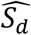 (adjusted for 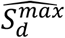 and 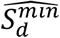) averaged over all FOV dimensions. The results are given in **Table 1** and show that the MSSL of isogenic and kin strains is the same (within experimental error), but this differs from the MSSL of non-kin strains.

**FIG 2.**
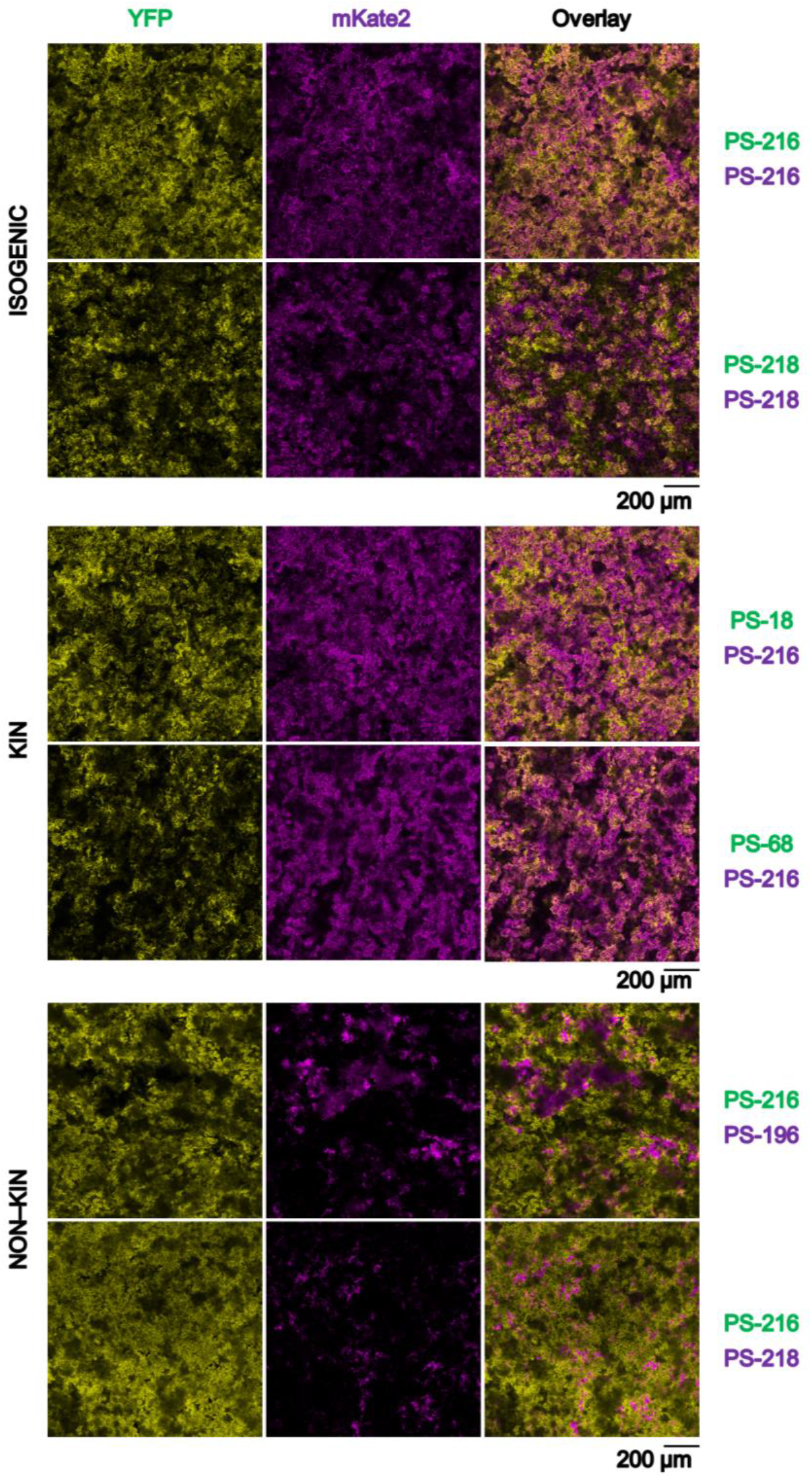
Segregation analysis of isogenic, kin and non-kin strains in biofilms. Pellicles formed by mixing of two isogenic, kin and non-kin *B. subtilis* PS strains that constitutively express fluorescent reporters. Biofilms were examined by confocal laser scanning microscopy (scale bar, 200 μm) after 16 hours of biofilm growth. This time point was chosen to ensure imaging before sporulation started. Images were pseudo-colored yellow (for YFP tagged *B. subtilis* strains PS-216, PS-218, PS-18, PS-68) PS-) and magenta (for mKate2 tagged strains *B. subtilis* PS-216, PS-218, PS-196). Representative images of at least three biological replicates are shown.

**FIG 3.**
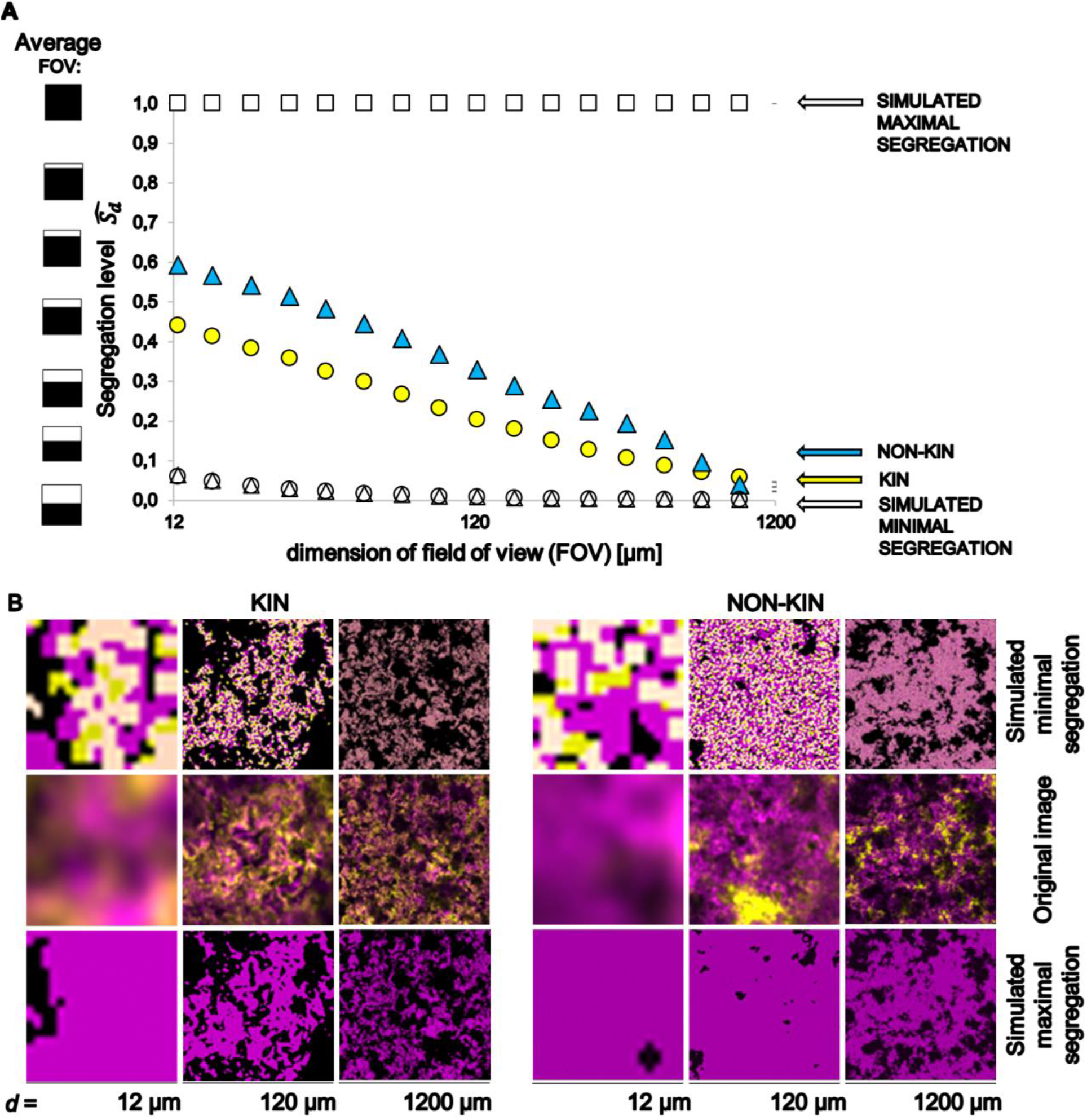
Multiscale spatial segregation analysis (MSSA) of biofilm image stacks obtained by confocal laser scanning fluorescent microscopy (CLSM). (A) Plots of calculated segregation levels 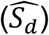 in mixed strain biofilms are shown for the kin pair (PS-68 YFP + PS-216 mKate2) strains and non-kin pair (PS-218 YFP + PS-216 mKate2) strains as a function of the dimension of field of view (FOV). Examined FOV, which represents what is seen under the microscope at different zoom levels, is a square with the dimensions *d* x *d*. 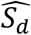 was calculated for simulated maximal and minimal segregations. 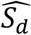 is 1 if only one strain is present in the FOV and 0 if both strains are present in the expected ratios (ratio of the two strains in all images of the stack) in the FOV, graphically represented as the average FOV. (B) Microscopy images (middle slice of CLSM stack) representing the FOV with *d* of 12 μm, 120 μm and 1200 μm. Top and bottom rows show corresponding simulated images; middle row represents original images as taken by the microscope. For the simulation of maximal segregation, it was assumed that one strain completely excluded the other strain.

**TABLE 1:** Multiscale spatial segregation levels (MSSLs) of isogenic, kin and non-kin two strain biofilms ^a^.

| A MSSL values for each strain combinations |  |  |  |  |  |
| --- | --- | --- | --- | --- | --- |
|  | PS-18 | PS-68 | PS-196 | PS-216 | PS-218 |
| PS-18 | 0.10 |  |  |  |  |
| PS-68 |  | 0.12 |  |  | 0.20 |
| PS-196 |  |  | 0.11 | 0.21 |  |
| PS-216 | 0.12 | 0.09 | 0.25 | 0.10 | 0.14 |
| PS-218 |  | 0.17 |  | 0.21 | 0.10 |
| B |  |  |  |  |  |
|  | ISOGENIC | KIN | NON-KIN |  |  |
| AVG | 0.10 | 0.11 | 0.21 |  |  |
| SD | 0.01 | 0.02 | 0.03 |  |  |

<sup>a</sup> MSSLs values for the tested strain combinations (A). Average (AVG) and standard deviation (SD) of MSSL values for the particular type of strain combination (B). Purple and green numbers represent the *B. subtilis* PS strains labelled by mKate2 and YFP, respectively. Grey fields denote isogenic combinations, yellow kin combinations and blue fields non-kin combinations. Larger MSSL value indicates more intense segregation.

### KD limits EPS sharing in conditions favoring matrix producers

The extracellular matrix is considered a public good, which is shared by neighboring cells but which could be exploited by “mutants” that benefit from the cooperation without paying the cost (1, 33). *B. subtilis* 3610 wild-type (WT), which forms a robust pellicle, shares its EPS with the Δ*epsA-O* mutant and hence complements the mutant defect in forming a floating pellicle (26, 28, 34). However, we have limited insights into the matrix sharing between strains from different KD groups, which we address here. A recent report indicates that the overall fitness of the matrix mutants remained stable even when the visible pellicle did not form due to the lack of matrix components (35). Therefore, we aimed to develop an experimental set-up in which the matrix non-producer (Δ*epsA-O*) will show a fitness disadvantage compared to the WT matrix producer. We assumed that by increasing the volume of the media we would increase the oxygen gradient, and hence provide the advantage to the WT over the mutant, which will have a disadvantage in invading the oxygen-replete surface–air interface. To identify the desired conditions, we grew the WT PS-216 and its mutant in 4, 20 and 50 ml media in 50-ml tubes covered with loosened lids. Although the pellicle of the PS-216 Δ*epsA-O* mutant was very weak it was still formed at the liquid surface even in 50 ml volume (**Fig. S3A)**. To provide a fair fitness comparison for all tested strains we counted the overall number of spores in sonicated biofilms and in the underlying medium (in total volume) in monocultures of the WT and the mutant after 72 hours of static incubation using phase-contrast microscopy (**Fig. S3B**). The spore counts in monocultures of the PS-216 Δ*epsA-O* were comparable to the WT counts in 20 ml and 4 ml volumes but decreased for 0.66-log CFU/ml (*P* = 0.046) compared to the WT in 50 ml volume. Hence, the mutant had a fitness disadvantage only in larger media volume, where the selective pressure on strains that fail to efficiently occupy the oxygen-rich surface should be more pronounced. Next, we compared PS-216 and PS-196 WT strains and their respective Δ*epsA-O* mutants in monocultures grown in a 50 ml medium with a loosely closed lid (**Fig. 4**). In this experiment we counted the CFU only in pellicles, where PS-196 had a significant fitness advantage compared to its respective Δ*epsA-O* mutant, which reached about 3.34-log CFU/ml (*P* = 0.001) lower counts than the parental WT. Although PS-216 WT showed a lesser advantage over the mutant PS-216 Δ*epsA-O* (0.8 log CFU/ml, *P* = 0.001) in pellicles (**Fig. 4**) than the PS-196 wild type over its matrix mutant, these experiments confirmed that the 50 ml volume set-up with loosened lids ensures conditions in which WT has a fitness advantage over the Δ*epsA-O* mutant. Therefore, this experimental set-up is appropriate for testing matrix sharing between the WT and the mutant in floating biofilms. We predicted that sharing will be more efficient in kin mixes than in non-kin mixes. To test this, we mixed the Δ*epsA-O* with the parental WT strain (at a 1:1 starting ratio). PS-196 efficiently complemented the mutant phenotype in co-culture and the mutant increased significantly (*P* = 0.002) in CFU counts (3-log) compared to its monoculture. However, the PS-216 Δ*epsA-O* mutant already had high fitness in the monoculture, and we did not observe a further increase in the co-culture **(Fig. 4)**, which suggests that the production of EpsA-O is less important for pellicle formation in this strain.

**FIG 4.**
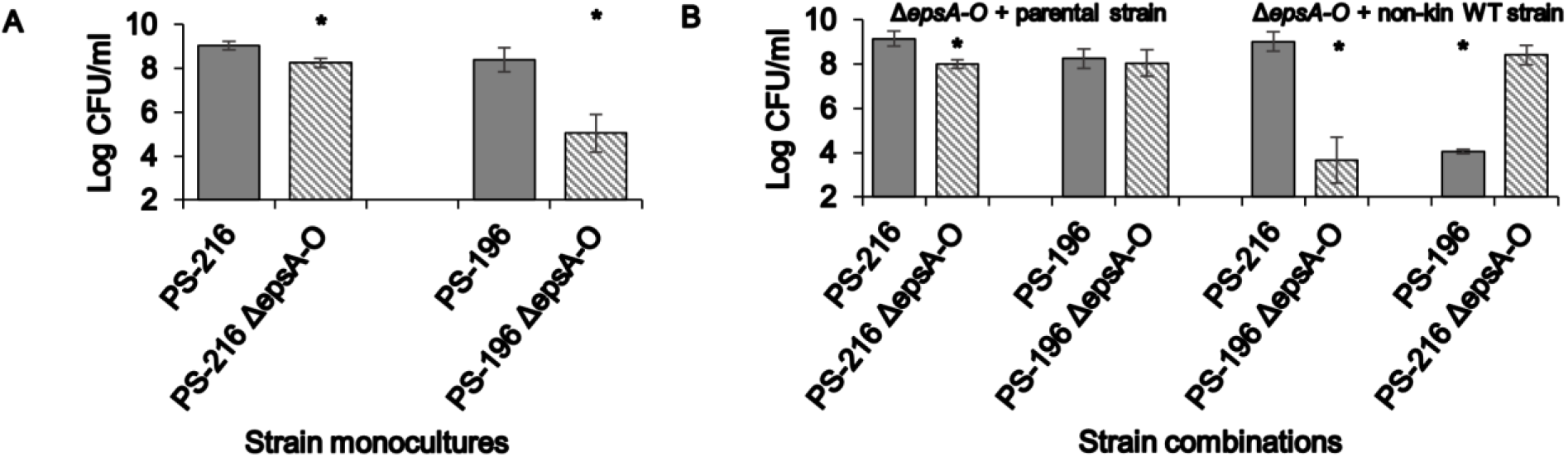
A WT producer complements kin strains but not non-kin strains with a polysaccharide matrix defect (*Δeps*) in mixed pellicles. Fitness of each strain grown alone (a), or in a pellicle co-culture with the *Δeps* mutant paired with the parental or non-kin WT (b), was measured by CFU counts after 24 hours of growth; experiments were performed in at least three to four independent experimental variants (n ≥ 3). *Indicates that the log CFU/ml is significantly different between the two strains monocultures or between the strain in mixed pellicles compared to its monoculture (*t*-Student; two-tail * *P* < 0.05).

Finally, we grew mixed biofilms composed of a non-kin matrix producer and non-producer. Here, PS-216 always outcompeted PS-196, regardless of whether it produced EPS or not, and was not affected by PS-196 itself, even in 50 ml cultures. The average fitness of the PS-196 matrix producer paired with the non-kin PS-216 Δ*epsA-O* mutant decreased dramatically (4.32-log CFU/ml; *P* = 0.001) compared to the CFU counts achieved when PS-196 was grown alone (**Fig. 4**). Overall, these results suggest that matrix was freely shared between the parental producer PS-196 and PS-196 *Δeps*, which is consistent with previous results (26, 28, 34), but not if PS-196 *Δeps* was paired with non-kin PS-216 matrix producer. The observed effect is most likely due to the exclusion of PS-196 by the dominant PS-216 strain (**Fig. 1)**.

## Discussion

Biofilms are composed of matrix-embedded bacteria that are capable of various social interactions that might affect fitness, cell assortment and matrix sharing, and consequently structure and ecology of microbial communities (9). By using a subset of microscale soil strains with previously determined kin types (13), we here confirm antagonism in non-kin in floating biofilms, while kin and isogenic strains coexist more peacefully. Non-kin also segregate into patches of different size and less freely share their matrix than kin.

We find a significant reduction in CFU counts of one strain in mixed biofilms composed of two non-kin strains, with one strain showing a significant dominance over the other. For example, PS-216 won over PS-196 or PS-218. In contrast, the fitness of both strains in kin and isogenic biofilms was preserved. The partial competitive exclusion of one strain in non-kin biofilms may be a consequence of antagonism between non-kin, which has been observed at the meeting point of two non-kin swarms of *B. subtilis* (13, 16, 18), *Proteus mirabilis* (36, 37) and *Myxococcus xanthus* (14, 38). Our results are also similar to those of previous studies on the consequences of mixing genetically divergent *Pseudomonas aeruginosa* strains in static cultures (39, 40). Although these authors did not specifically address KD, they reported antagonism between genetically divergent strains in biofilms. It is generally believed that clonemates cooperate, whereas genetically different cell lineages of the same species compete (1, 41, 42). Competitive strategies between non-kin comprise contact-dependent killing (43) and local exchange of antagonistic molecules between swarms of *B. subtilis* (16), *P. mirabilis* (12) and *M. xanthus* (44). Although swarms represent a different type of collective than pellicles, it is possible that non-kin deploy similar types of interactions during swarming and pellicle development, although there are also pronounced differences in the opportunities for social interactions. During a swarming encounter assay, cells in the swarm are initially surrounded by their kin and only at the point of encounter of two non-kin swarm cells do they engage in direct cell-cell contact. At this point they are exposed to higher cell density and nutrient/space limitation. During pellicle formation, non-kin first engage in interactions in the liquid medium, where they exist as planktonic cells, interacting through secreted and diffusible factors and even extracellular polymers that engage cells in a loose network (5). Several hours later, probably when they sense a limitation of oxygen, they invade the air–liquid interface, start forming patches and engage in direct cell-cell interactions (30). Our results demonstrate that in pellicles bacterial cells segregate into visible patches similar to those that have previously been shown for colonies (45, 46). However, the assortment of non-kin cells produces larger patches of the dominant strain that are interspersed by smaller patches of the less competitive non-kin strain. This is different from assortment in kin pellicles, where mixing is more pronounced and less heterogenous. This assortment is consistent with modeling experiments testing inter-strain competition in spatially structured environments between a bacteriocin producer and non-producer, which resulted in killing-driven assortment of two genotypes (47, 48).

In addition to social interactions, the biofilm heterogeneity is also shaped by steep vertical oxygen gradients that result from oxygen consumption by bacteria and slow oxygen diffusion into the medium (5, 49, 50). Pellicle construction is a strategy for effectively obtaining oxygen at the liquid surface (24, 25, 51). Therefore, in our experimental setting with high medium volume, and thus more substantial selective pressure to swim toward the surface to reach the environment with available oxygen, EPS producers (PS-216 and PS-196) had a fitness advantage in pellicles over the EPS non-producers (PS-216 Δ*epsA-O*, PS-196 Δ*epsA-O*), and even in total volume at the level of spore counts. A possible explanation for the role of matrix in maintaining pellicles at the liquid surface is a decrease in the specific gravity or/and intrinsic hydrophobicity of cells glued by a matrix, which counteracts selective forces faced by planktonic cells (51). We observed that the benefit of EPS for occupying the surface was more important for the PS-196 than the PS-216, which even without EPS still formed a weak pellicle at the liquid surface, and we observed that the decrease in cell counts for PS-216 mutant compared to its WT was much lower than that detected for the PS-196 mutant strain (**Fig. 4**). PS-216 may encode additional polysaccharides that compensate for the lack of EpsA-O in the mutant. In fact, natural isolates of *B. subtilis* often carry the intact copy of the *ypqP* (*spsM*) gene, encoding a sugar epimerase likely to be involved in polysaccharide synthesis (52), which is inactivated due to the integration of the SPβ prophage in the standard biofilm NCIB3610 strain and the laboratory strain 196 (53). The *spsM* gene is also intact in the PS-216 genome (18). Alternatively, PS-216 is well adapted to oxygen deficiency and therefore its growth may be less affected by oxygen gradient, as we have shown recently (54) and therefore overall fitness of the strain is less sensitive to the lack of the EPS polysaccharide.

We also studied matrix sharing between the Δ*epsA-O*-mutant and the parental or non-kin producer at the air–liquid interface. The PS-196 Δ*epsA-O* mutant was able to utilize Eps A-O in pellicle produced by its parental strain, confirming previous work performed with NCBI3610 strain (26, 28, 34). In contrast, the PS-216 Δ*epsA-O* mutant did not gain any advantage from the interaction with the parental strain and its fitness remained similar to that in the monoculture. Privatization of the PS-216 EpsA-O matrix was especially prominent in the co-culture with the PS-196 Δ*epsA-O* mutant, whose fitness decreased 10,000-fold compared to its fitness in co-culture with PS196 WT and did not improve compared to the fitness in mutant monoculture. Also, fitness of the PS-196 WT in co-culture with the PS-216 Δ*epsA-O* decreased significantly. This suggests that PS-216 antagonizes the PS-196 strain, and that through this mechanism KD limits matrix sharing between *B. subtilis* strains.

Additional studies have investigated possible strategies to limit matrix sharing in mixed pellicles composed of matrix producers and non-producers that are kin in other loci. For example, Irie *et al*. (2017) observed that the distribution of matrix in *P. aeruginosa* biofilms is social and is shared with other producers (55), but non-producers were unable to utilize it, suggesting that matrix could only be shared locally with surrounding cells that are usually genetically identical, which assured high relatedness in a local patch (55). Moreover, in pellicles of S*almonella enterica* serovar a matrix component such as cellulose kept non-producers away from the producers (56), which is consistent with the observation of limited matrix sharing between a PS-216 producer and non-producer, where some other matrix component might potentially interfere with matrix sharing.

In conclusion, our results support the hypothesis that *B. subtilis* strains harbor antagonistic mechanisms directed toward non-kin strains, which operate through a fitness reduction of one strain and a modified assortment of non-kin groups in the floating biofilm. These mechanisms may limit the invasion of the WT by non-kin matrix non-producers, and thereby biofilm matrix exploitation.

## MATERIALS AND METHODS

### Strains and media

Strains used in this study are described in **Table S1** Briefly, WT *Bacillus subtilis* strains were isolated two samples of 1 cm3 of soil from the sandy bank of the Sava River in Slovenia previously (32). Recombinant strains carried reporters for YFP or red fluorescent protein (mKate2) from constitutively expressed promoters. Strains PS-216-Phyperspank-mKate2 and PS-218-Phyperspank-mKate2 were constructed previously (13) or were created using a standard protocol by transforming the chromosomal DNA from a known recombinant strain into the *amyE* locus by homologous recombination and selection for antibiotic resistance – chloramphenicol (Cm) 5 μg/ml for strains tagged with mKate2 or spectinomycin (Sp) 100 μg/ml for YFP tagging.

The *B. subtilis* strains WT PS-216 Δ*epsA-O*::tet *amyE*::P_*hyperspank*_-*mKate2* and PS-196 Δ*epsA-O*::tet *amyE*::P_*hyperspank*_-*mKate2* were obtained via transformation of the strains PS-216 P_*hyperspank*_-*mKate2* (13) and PS-196 P_*hyperspank*_-*mKate2*, respectively, with genomic DNA originating from the strain *B. subtilis* NCIB 3610 Δ*epsA-O*::tet (21) and selected for antibiotic resistance to tetracycline (Tet) 10 μg/ml.

Strains with constitutively expressed fluorescent protein (mKate2) were PS-196-P_*hyperspank*_-*mKate2*, PS-68-P_*hyperspank*_-*mKate2*, PS-18-P_*hyperspank*_-*mKate2*, which were constructed by transforming the WT PS-196, PS-68 and PS-18 competent cells, respectively, with the chromosomal DNA isolated from the strain *B. subtilis* NCIB 3610 CY49 and selecting transformants obtained by double cross-over on antibiotic chloramphenicol containing LB agar (Table S1).

Strains PS-216-P_*hypercIo3*_-*yfp*, PS-196-P_*hypercIo3*_-*yfp*, PS-218-P_*hypercIo3*_-*yfp*, PS-68-P*_hypercIo3_*-*yfp*, and PS-18-P_*hypercIo3*_-*yfp* were obtained by transforming the WT PS-216, PS-196, PS-218, PS-68 and PS-18, respectively, with chromosomal DNA isolated from the strain *B. subtilis* NCIB 3610 P_*hypercIo3*_-*yfp* (a kind gift from Roberto Kolter) (16) as described in Table S1.

Strains were routinely grown in LB medium. MSgg medium was used for biofilm growth and was prepared by combining 20.93 g MOPS, 1.06 g K3PO4, 0.05 g tryptophan, 0.05 g phenylalanine, 0.41 g MgCl2 x 6H20, 5 g Na-glutamate, and 5 g glycerol, supplemented with 1 ml of microelements (700 μM CaCl2, 50 μM FeCl3, 50 μM MnCl2, 1 μM ZnCl2) per 1 l of dH2O. pH value was adjusted to 7. After autoclaving, just before performing an experiment, 2 μM (1 ml/l) of thiamine hydrochloride was added (24). LB medium for agar plates was prepared by adding 35 g of LB agar (Lennox) per 1 l of dH2O, solidified through the addition of agar (Sigma-Aldrich) to 2%. The plates were allowed to dry at room temperature overnight.

### Biofilm competition assays

Biofilm competition assays were performed in liquid MSgg medium, as previously described (24). Inoculum of each strain was prepared from spores revitalized in LB for 2 hours (shaking at 200 rpm at 37°C) until they reached an optical density (O. D.) of around 0.2. Two strains were inoculated at a 1:1 ratio (10^5^ cells/ml each). Initial colony count (CFU) assays were performed immediately after inoculation on LB agar plates containing spectinomycin (100 μg/ml) for strains labeled with YFP or chloramphenicol (5 for μg/ml) for strains with the mKate2 fluorescence marker. Biofilms were grown as static cultures, which promoted the formation of pellicles at the liquid–air interphase, in 2 ml of MSgg medium in a 12-well plate at 37°C for 24 hours. After incubation, pellicles were harvested by pipette tips, transferred to 1 ml of physiological saline and disintegrated by ultrasound sonicator (MSE 150 Watt Ultrasonic Disintegrator Mk2: 3 x 10 seconds with 20 seconds of pause between each cycle at 20kHz with an amplitude of 15 μm) to obtain single cells for CFU determination. Sonicated pellicles were serially diluted (by 10-fold steps) in physiological saline and plated on LB agar plates containing the same antibiotics as for initial CFU determination.

### Microscopy

To examine the spatial distribution of two strains, we again used *B. subtilis* strains labeled with constitutively expressed YFP or mKate2. Mixed cultures were grown in 2 ml of MSgg medium in 12-well plates as described above (see **Competition biofilm assays**). After 16 hours of incubation at 37°C, before sporulation started, the medium below the pellicle was removed and pellicles were subjected to CLSM Axiovsion Z1, LSM800 (Zeiss, Germany). A16 hour time point was chosen because at that time sporulation has not yet been initiated. The CLSM images were acquired by EC Plan-Neofluar 10x/0.30 Ph 1 objective by two laser channels: the 488 nm laser to acquire fluorescence from the YFP protein and the 561 nm laser to acquire fluorescence from the mKate2 protein. The emitted fluorescence was recorded at 400–600 nm and 600–700 nm for YFP and mKate2, respectively. Pinhole size for YFP was set to 1.0 AU and for mKate2 it was set to 1.2 AU. Frame time was 3.7 seconds, averaging 4 seconds, and pixel single frame size was 930 x 930. The mosaic function of Zen 2.3 (Zeiss, Germany) was used for 2 x 2 frame acquisition. The stitched image of 1960 x 1960 pixels covered a FOV with dimensions 1.2 mm x 1.2 mm. Slicing was set to half of the Nyquist distance, which was 6 μm.

To improve the quality and resolution, the CLSM images were deconvolved by Tikhonov-Miller algorithm applied in DeconvolutionLab2 application (57) with artificial point spread function as obtained by PSF generator application (58), both running in a Fiji-ImageJ environment (59). Images were then manually thresholded and converted into binary format, which served as an input for MSSA using the recently developed ImageJ tool (for details see Dogsa *et al*. (submitted).

### Multiscale spatial segregation analysis (MSSA) of CLSM biofilm images

MSSA determines the segregation level among two strains as a function of the size of the space (FOV of dimensions *d* x *d*) that two strains co-occupy in a biofilm (detailed information available in Dogsa *et al*., submitted). The *d_min_* corresponded to two times the Nyquist distance (12 μm) and *d_max_* to 1.2 mm. The FOV of different *d* is randomly placed on an image and the strain abundance is presented as area normalized to the average abundance of the two strains. We calculated the local segregation level (i.e. the segregation in the specific FOV) as the relative difference in the abundance of one strain compared to the other. By weight averaging many local segregation levels and changing the dimension, *d* of FOV from *d_min_* to *d_max_*, we determined the segregation level, 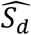 as a function of *d*. The segregation level ranges from 0 to 1, where 0 represents no segregation (two differentially labeled strains are simultaneously present in the same FOV in expected amounts) and 1 represents ideal segregation (two differentially labeled cells are never found simultaneously in the examined FOV). The exact values of two extremes of segregation levels, denoted as 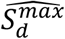 and 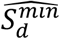, are, however, dependent on a particular image and experimental conditions and can be simulated by our approach.

The sampling factor was typically 0.005, which means that around 10,000 randomly placed FOVs were evaluated at *d_min_*. The standard error of the segregation level was in all dimensions of FOV < 2%. The resolution factor, which determines the spacing among FOV dimensions for which segregation level will be determined, was set to 0.75. This means that from *d_max_* to *d_min_* of FOV we have 22 points on the graph (**Fig. 3A**), which represents the segregation level vs dimension (*d*) of FOV (distances between points between *d_max_* to *d_min_* decrease by the factor 0.75). To simulate the minimal segregation, the bacterial cell was assumed to be a circle of 1.8 μm in diameter, which corresponded well to sizes determined in sonicated samples, where we could resolve the size of the individual cells. During simulations the cells were randomly distributed in biofilm of the same shape as experimental biofilms and overlap between cells was allowed. For the calculation of the maximal segregation level, it was assumed that one strain completely excluded the other strain and thus only this strain is present in each image slice. Also, it was further assumed that the shape of the original biofilm is preserved.

### Experiments testing the effect of oxygen limitation on fitness of the WT and the Δ*epsA-O* mutant in co-cultures

The effect of oxygen availability on biofilm growth of *B. subtilis* PS-216 and PS-216 Δ*epsA-O* mutant was examined in 50 ml and also in 20 and 4ml of MSgg medium in Falcon tubes with loosened lids (higher oxygen concentration at the liquid surface). Each tube was inoculated by approximately 10^5^ cells/ml and pellicles were grown at static conditions for 24 hours at 37°C (for details see figure legend of Figure S3).

Further experiments addressing competition between the WT and the Δ*epsA-O*-mutant strains with the parental and non-kin combination were investigated in 50 ml of MSgg medium in Falcon tubes with loosened lids. Inocula were prepared from overnight cultures of each strain grown in 5 ml of LB medium with spectinomycin (100 μg/ml) or chloramphenicol (5 for μg/ml) and tetracycline (10 μg/ml) for 16 – 18 hours (shaking at 200 rpm at 37°C) and then 100 x diluted in 5 ml of fresh LB medium with the same antibiotics. Shaken cultures were grown for an additional 2.5 hours to mid-log phase and then re-inoculated in 5 ml of LB for another 2.5 hours to reach an O. D. value of around 0.2. This ensured comparable physiological states of each strain used as an inoculum to set up the mixed culture experiments of two strains. After incubation for 24 hours at 37°C at static conditions, the CFU assay was performed as described above (see **Competition biofilm assays**).

### Statistical analysis

All experiments were performed in at least three biologically independent replicates. Bacterial counts are log_10_-transformed and data are expressed as average values ± standard deviations of the means. Statistical differences between the CFU of the strains in mono/co-cultures were determined using a two-sample Student’s *t*-test. *P*-values of less than 0.05 were considered significant.

## ACKNOWLEDGMENTS

This research has been financially supported by the Slovenian Research Agency (ARRS) program grant P4-0116, and by project grant J4-9302 (awarded to I. Mandic Mulec). We also acknowledge the Microscopy of Biological Samples infrastructural center, located in the Biotechnical Faculty at the University of Ljubljana, Slovenia. We thank Tjaša Stošicki for her help with the initial experiments. We also thank Polonca Štefanič and Anna Dragoš for helpful discussions and Roberto Kolter for kind gift of NCB3610 strains.

## References

1. Nadell CD, Drescher K, Foster KR. 2016. Spatial structure, cooperation and competition in biofilms. Nat Rev Microbiol 14:589–600.

2. Elias S, Banin E. 2012. Multi-species biofilms: living with friendly neighbors. FEMS Microbiol Rev 36:990–1004.

3. Flemming HC, Wuertz S. 2019. Bacteria and archaea on Earth and their abundance in biofilms. Nat Rev Microbiol 17:247–260.

4. Burmolle M, Ren D, Bjarnsholt T, Sorensen SJ. 2014. Interactions in multispecies biofilms: do they actually matter? Trends Microbiol 22:84–91.

5. Flemming HC, Wingender J, Szewzyk U, Steinberg P, Rice SA, Kjelleberg S. 2016. Biofilms: an emergent form of bacterial life. Nat Rev Microbiol 14:563–75.

6. Drescher K, Nadell CD, Stone HA, Wingreen NS, Bassler BL. 2014. Solutions to the public goods dilemma in bacterial biofilms. Curr Biol 24:50–55.

7. Hamilton WD. 1964. The genetical evolution of social behaviour. I. J Theor Biol 7:1–16.

8. Hamilton WD. 1964. The genetical evolution of social behaviour. II. J Theor Biol 7:17–52.

9. Gilbert OM, Strassmann JE, Queller DC. 2012. High relatedness in a social amoeba: the role of kin-discriminatory segregation. Proc Biol Sci 279:2619–24.

10. Strassmann JE, Gilbert OM, Queller DC. 2011. Kin discrimination and cooperation in microbes. Annu Rev Microbiol 65:349–67.

11. Dienes L. 1946. Reproductive processes in Proteus cultures. Proc Soc Exp Biol Med 63:265–70.

12. Alteri CJ, Himpsl SD, Pickens SR, Lindner JR, Zora JS, Miller JE, Arno PD, Straight SW, Mobley HL. 2013. Multicellular bacteria deploy the type VI secretion system to preemptively strike neighboring cells. PLoS Pathog 9:e1003608.

13. Stefanic P, Kraigher B, Lyons NA, Kolter R, Mandic-Mulec I. 2015. Kin discrimination between sympatric Bacillus subtilis isolates. Proc Natl Acad Sci U S A 112:14042–7.

14. Vos M, Velicer GJ. 2009. Social conflict in centimeter-and global-scale populations of the bacterium Myxococcus xanthus. Curr Biol 19:1763–7.

15. Wielgoss S, Fiegna F, Rendueles O, Yu YN, Velicer GJ. 2018. Kin discrimination and outer membrane exchange in Myxococcus xanthus: A comparative analysis among natural isolates. Mol Ecol 27:3146–3158.

16. Lyons NA, Kraigher B, Stefanic P, Mandic-Mulec I, Kolter R. 2016. A Combinatorial Kin Discrimination System in Bacillus subtilis. Curr Biol 26:733–42.

17. Jain C, Rodriguez RL, Phillippy AM, Konstantinidis KT, Aluru S. 2018. High throughput ANI analysis of 90K prokaryotic genomes reveals clear species boundaries. Nat Commun 9:5114.

18. Stefanic P, Belcijan K, Kraigher B, Kostanjšek R, Nesme J, Madsen J, Kovac J, Sørensen S, Vos M, Mandic-Mulec I. 2021. Kin discrimination promotes horizontal gene transfer between unrelated strains in Bacillus subtilis. Nature commun, in press doi:10.1101/756569.

19. Kalamara M, Spacapan M, Mandic-Mulec I, Stanley-Wall NR. 2018. Social behaviours by Bacillus subtilis: quorum sensing, kin discrimination and beyond. Mol Microbiol 110:863–878.

20. Bais HP, Fall R, Vivanco JM. 2004. Biocontrol of Bacillus subtilis against infection of Arabidopsis roots by Pseudomonas syringae is facilitated by biofilm formation and surfactin production. Plant Physiol 134:307–19.

21. Beauregard PB, Chai Y, Vlamakis H, Losick R, Kolter R. 2013. Bacillus subtilis biofilm induction by plant polysaccharides. Proc Natl Acad Sci U S A 110:E1621–30.

22. Verhamme DT, Murray EJ, Stanley-Wall NR. 2009. DegU and Spo0A jointly control transcription of two loci required for complex colony development by Bacillus subtilis. J Bacteriol 191:100–8.

23. Vlamakis H, Chai Y, Beauregard P, Losick R, Kolter R. 2013. Sticking together: building a biofilm the Bacillus subtilis way. Nat Rev Microbiol 11:157–68.

24. Branda SS, Gonzalez-Pastor JE, Ben-Yehuda S, Losick R, Kolter R. 2001. Fruiting body formation by Bacillus subtilis. Proc Natl Acad Sci U S A 98:11621–6.

25. Holscher T, Bartels B, Lin YC, Gallegos-Monterrosa R, Price-Whelan A, Kolter R, Dietrich LEP, Kovacs AT. 2015. Motility, Chemotaxis and Aerotaxis Contribute to Competitiveness during Bacterial Pellicle Biofilm Development. J Mol Biol 427:3695–3708.

26. Branda SS, Chu F, Kearns DB, Losick R, Kolter R. 2006. A major protein component of the Bacillus subtilis biofilm matrix. Mol Microbiol 59:1229–38.

27. Romero D, Aguilar C, Losick R, Kolter R. 2010. Amyloid fibers provide structural integrity to Bacillus subtilis biofilms. Proc Natl Acad Sci U S A 107:2230–4.

28. Romero D, Vlamakis H, Losick R, Kolter R. 2011. An accessory protein required for anchoring and assembly of amyloid fibres in B. subtilis biofilms. Mol Microbiol 80:1155–68.

29. Branda SS, Gonzalez-Pastor JE, Dervyn E, Ehrlich SD, Losick R, Kolter R. 2004. Genes involved in formation of structured multicellular communities by Bacillus subtilis. J Bacteriol 186:3970–9.

30. Dragos A, Kiesewalter H, Martin M, Hsu CY, Hartmann R, Wechsler T, Eriksen C, Brix S, Drescher K, Stanley-Wall N, Kummerli R, Kovacs AT. 2018. Division of Labor during Biofilm Matrix Production. Curr Biol 28:1903–1913 e5.

31. Dragos A, Martin M, Falcon Garcia C, Kricks L, Pausch P, Heimerl T, Balint B, Maroti G, Bange G, Lopez D, Lieleg O, Kovacs AT. 2018. Collapse of genetic division of labour and evolution of autonomy in pellicle biofilms. Nat Microbiol 3:1451–1460.

32. Stefanic P, Mandic-Mulec I. 2009. Social Interactions and Distribution of Bacillus subtilis pherotypes at microscale. Journal of Bacteriology 191:1756–1764.

33. Nadell CD, Xavier JB, Foster KR. 2009. The sociobiology of biofilms. FEMS Microbiol Rev 33:206–24.

34. Martin M, Dragos A, Holscher T, Maroti G, Balint B, Westermann M, Kovacs AT. 2017. De novo evolved interference competition promotes the spread of biofilm defectors. Nat Commun 8:15127.

35. Steinberg N, Rosenberg G, Keren-Paz A, Kolodkin-Gal I. 2018. Collective Vortex-Like Movement of Bacillus subtilis Facilitates the Generation of Floating Biofilms. Front Microbiol 9:590.

36. Gibbs KA, Urbanowski ML, Greenberg EP. 2008. Genetic determinants of self identity and social recognition in bacteria. Science 321:256–9.

37. Gibbs KA, Greenberg EP. 2011. Territoriality in Proteus: advertisement and aggression. Chem Rev 111:188–94.

38. Gong Y, Zhang Z, Zhou XW, Anwar MN, Hu XZ, Li ZS, Chen XJ, Li YZ. 2018. Competitive Interactions Between Incompatible Mutants of the Social Bacterium Myxococcus xanthus DK1622. Front Microbiol 9:1200.

39. Oluyombo O, Penfold CN, Diggle SP. 2019. Competition in Biofilms between Cystic Fibrosis Isolates of Pseudomonas aeruginosa Is Shaped by R-Pyocins. mBio 10.

40. Oliveira NM, Martinez-Garcia E, Xavier J, Durham WM, Kolter R, Kim W, Foster KR. 2015. Biofilm Formation As a Response to Ecological Competition. PLoS Biol 13:e1002191.

41. Nadell CD, Foster KR, Xavier JB. 2010. Emergence of spatial structure in cell groups and the evolution of cooperation. PLoS Comput Biol 6:e1000716.

42. Mitri S, Foster KR. 2013. The genotypic view of social interactions in microbial communities. Annu Rev Genet 47:247–73.

43. Aoki SK, Pamma R, Hernday AD, Bickham JE, Braaten BA, Low DA. 2005. Contact-dependent inhibition of growth in Escherichia coli. Science 309:1245–8.

44. Dey A, Vassallo CN, Conklin AC, Pathak DT, Troselj V, Wall D. 2016. Sibling Rivalry in Myxococcus xanthus Is Mediated by Kin Recognition and a Polyploid Prophage. J Bacteriol 198:994–1004.

45. van Gestel J, Weissing FJ, Kuipers OP, Kovacs AT. 2014. Density of founder cells affects spatial pattern formation and cooperation in Bacillus subtilis biofilms. ISME J 8:2069–79.

46. Holscher T, Dragos A, Gallegos-Monterrosa R, Martin M, Mhatre E, Richter A, Kovacs AT. 2016. Monitoring Spatial Segregation in Surface Colonizing Microbial Populations. J Vis Exp doi:10.3791/54752.

47. Tait K, Sutherland IW. 2002. Antagonistic interactions amongst bacteriocin-producing enteric bacteria in dual species biofilms. J Appl Microbiol 93:345–52.

48. Bucci V, Nadell CD, Xavier JB. 2011. The evolution of bacteriocin production in bacterial biofilms. Am Nat 178:E162–73.

49. Stewart PS, Franklin MJ. 2008. Physiological heterogeneity in biofilms. Nat Rev Microbiol 6:199–210.

50. Werner E, Roe F, Bugnicourt A, Franklin MJ, Heydorn A, Molin S, Pitts B, Stewart PS. 2004. Stratified growth in Pseudomonas aeruginosa biofilms. Appl Environ Microbiol 70:6188–96.

51. Yamamoto K, Arai H, Ishii M, Igarashi Y. 2011. Trade-off between oxygen and iron acquisition in bacterial cells at the air-liquid interface. FEMS Microbiol Ecol 77:83–94.

52. Abe K, Kawano Y, Iwamoto K, Arai K, Maruyama Y, Eichenberger P, Sato T. Developmentally-regulated excision of the SPβ prophage reconstitutes a gene required for spore envelope maturation in Bacillus subtilis.

53. Sanchez-Vizuete P, Le Coq D, Bridier A, Herry JM, Aymerich S, Briandet R. Identification of ypqP as a New Bacillus subtilis biofilm determinant that mediates the protection of Staphylococcus aureus against antimicrobial agents in mixed-species communities.

54. Erega A, Stefanic P, Dogsa I, Danevčič T, Simunovic K, Klančnik A, Smole Možina S, Mandic Mulec I. 2021. Bacillaene Mediates the Inhibitory Effect of Bacillus subtilis on Campylobacter jejuni Biofilms.

55. Irie Y, Roberts AEL, Kragh KN, Gordon VD, Hutchison J, Allen RJ, Melaugh G, Bjarnsholt T, West SA, Diggle SP. 2017. The Pseudomonas aeruginosa PSL Polysaccharide Is a Social but Noncheatable Trait in Biofilms. mBio 8.

56. Srinandan CS, Elango M, Gnanadhas DP, Chakravortty D. 2015. Infiltration of Matrix-Non-producers Weakens the Salmonella Biofilm and Impairs Its Antimicrobial Tolerance and Pathogenicity. Front Microbiol 6:1468.

57. Sage D, Donati L, Soulez F, Fortun D, Schmit G, Seitz A, Guiet R, Vonesch C, Unser M. 2017. DeconvolutionLab2: An open-source software for deconvolution microscopy. Methods 115:28–41.

58. Kirshner H, Aguet F, Sage D, Unser M. 2013. 3-D PSF fitting for fluorescence microscopy: implementation and localization application. J Microsc 249:13–25.

59. Schindelin J, Arganda-Carreras I, Frise E, Kaynig V, Longair M, Pietzsch T, Preibisch S, Rueden C, Saalfeld S, Schmid B, Tinevez JY, White DJ, Hartenstein V, Eliceiri K, Tomancak P, Cardona A. 2012. Fiji: an open-source platform for biological-image analysis. Nat Methods 9:676–82.

